# Long-lasting humoral immunity in Covid-19 infected patients at a University Hospital Clinic in Östergötland County Council during 2020-2021

**DOI:** 10.1101/2021.04.27.441589

**Authors:** M Azharuddin, D Aili, R Selegård, Sajjad Naeimipour, M Sunnerhagen, HK Patra, K S Sjöberg, K Niward, H Hanberger, Å Östholm-Balkhed, J Hinkula

## Abstract

Longitudinal serum samples and nasopharyngeal/nasal swab samples were collected from forty-eight individuals (median age 66yrs) with Covid-19 PCR-positive test results at Linköping University Hospital. Samples were collected from initial visit and for 6 months follow up. Presence of serum IgG and IgA against SARS-CoV-2 antigens (S1-spike, nucleocapsid and NSP3) were analyzed. Nasal swabs were tested for presence of IgA against the outer envelope S1 spike protein. Ninety-two percent of participants were seropositive against SARS-CoV-2 recombinant proteins at day 28 from study entry and all (100%) were seropositive from samples collected at 2 months or later. The most common antibody responses (both serum IgG, mainly IgG1 and IgA) were detected against the S1-spike protein and the nucleoprotein. In samples collected from nasal tissues considerably lower frequencies of IgA-positive reactivities were detected. Sixteen to 18 percent of study participants showed detectable IgA levels in nasal samples, except at day 60 when 36% of tested individuals showed presence of IgA against the S1-spike protein. The study suggests that the absolute majority of studied naturally infected Covid-19 patient in the Linkoping, Ostergotland health region develop over 6 months lasting detectable levels of serum IgG and IgA responses towards the SARS-CoV-2 S1-spike protein as well as against the nucleoprotein, but not against the non-structural protein 3.

## Introduction

One of the cornerstones of protective immune response against viral infections is the humoral immunity, systemically and at mucosal surfaces. In most viral vaccine trials, in the childhood vaccination programmes (1–3) as well as in adult vaccination the humoral immunity is evaluated, and most often correlated with disease severity reduction or prevention. For respiratory tract infections such as influenza and RSV this is well documented (4–6). It is thus very likely so also with the Covid-19 infectious virus, the betacoronavirus SARS CoV-2 (7-11, 12). And this is the reason for conducting this study. The aim of this study is to follow a significant size of hospitalized Covid-19 patients from hospital admission and follow up to determine the longevity of the humoral immunity in blood and respiratory tract.

## Materials and methods

### Samples

Patients admitted to the hospital from whom sampled nasal swabs were shown as PCR-positive for SARS-CoV-2 (age range 24 – 81 years, median age 66) were asked for blood samples as well as additional nasal nasopharyngeal swabs. Samples were collected at day of admission, at 2 weeks, 4 weeks, 2 months and 6 months during follow-up. Negative and positive controls were obtained from laboratory staff at Linköping University.

### Serology

Serum was collected, aliquoted and frozen until used. Oral and nasal swabs were collected with cotton pads and kept in 1 mL sterile saline solution stored frozen until used. SARS-CoV-2 S1-spike protein (Wuhan strain 2019) was used as soluble antigen on 96-well microwell plates (0,5 ug/mL PBS pH 7,2, Toronto University, SGU, Toronto ON, Canada / SinoBiologicals, Eschborn, Germany). SARS-CoV-2 nucleoproteins NC-(aa47-174 and aa267-364) and nsp3 (Toronto University, ON, Canada). Serum samples from Covid-19 patients and positive and negative controls were diluted in PBS-Tween 20 (0.05%) with 2,5% fat-free milk buffer. Serum dilutions and standard serum control samples was added to the coated plate wells and incubated 90 min. at 37°C. Conjugates against anti-human IgG-HRP (BioRad, Richmond, CA) or anti-human IgA-HRP (Nordic BioSite, Täby, Sweden) was added to separate wells with diluted serum samples and incubated 90 min. at 37°C. Finally, substrate 0.003% H_2_O_2_/o-phenylene diamine (Sigma-Aldrich, S:t Louis, MA. 0,4 mg/mL) was added and kept at room temperature for 30 min. before 2,5M H_2_SO_4_ was added as stop solution. Plates were read at OD490. Standard curve samples for determining AU antibody quantitation was used. Serum IgG subclasses were tested with isotype-specific murine anti-human IgG subclasses (Sigma-Aldrich, S:t Louis, MA).

Mucosal samples from Covid-19 patients and positive and negative controls were diluted with PBS-Triton X-100 5% with 2,5% fat-free dry-milk and added to the coated SARS-CoV-2 S1 spike plate wells and incubated o.n min. at RT. Conjugates against anti-human IgA-HRP (Nordic BioSite, Täby, Sweden) was added and incubated for 90 min. at 37°C. Substrate and plate absorbance reading was performed as described above. Cut off was determined by identification of reactivity towards the antigens used from ten negative control serum mean reactivity values multiplied with two.

### Statistical analysis

Mann-Whitney U test was performed to compare differences in antibody reactivity between sampling time points.

Ethical permission: The study was conducted according to the rules of the Declaration of Helsinki. Samples were collected after informed consent and ethical permission was obtained and approved by the Ethical Review board in Sweden. (Reference no: 2020-03888 and 2020-02080).

## Results

### Serology

In total, forty-eight patients with Covid-19 infection were included. The kinetics of serum IgG and IgA against the SARS-CoV-1 S1-spike protein was analyzed (Figure 1A and 1B).

**Figure 1.**
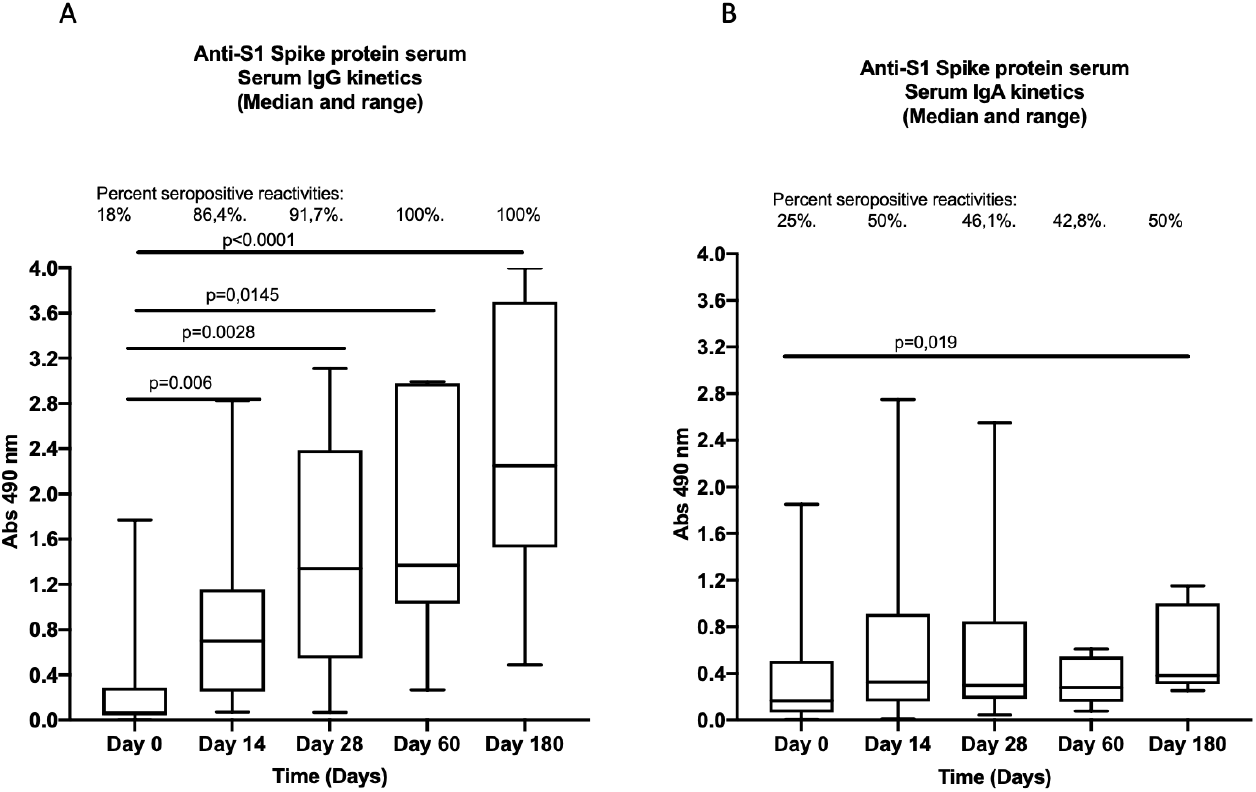
A. Kinetics of serum IgG reactivity against the SARS-CoV-2 S1 spike protein in forty-eight Covid-19 patient samples collected during 6 months from hospital admission. Boxplots showing median reactivity and range per timepoint is shown. B. Kinetics of serum IgA reactivity against the SARS-CoV-2 S1 spike protein in forty-eight Covid-19 patient samples collected during 6 months from hospital admission. Boxplots showing median reactivity and range per timepoint is shown.

All patients developed detectable IgG and IgA responses at least once during the study period. All of the study participants were seropositive at 1 month and remained serum IgG seropositive still at 6 months follow up. Serum IgA seropositivity was detectable in all tested samples at 2 months but was reduced to 78,6% at 6 months (not shown). Subclass IgG-isotyping was performed, and only IgG1 subclass IgG was detected.

Serum IgG against SARS-CoV-2 nucleoproteins (NC-protein aa 47-173) was seen with serum from all study participants at 1 month post admission. However, only 2 study participants responded with detectable serum IgG against a more C-terminal part of the nucleoprotein (aa 247-364). The comparison of anti-NC-binding antibodies in serum was compared with the serum IgG anti-S1 spike reactivity (Figure 2). Serum IgG and IgA reactivity against one non-structural protein, the NSP3-protein was assayed, and all tested sera were shown to be non-reactive (not shown).

**Figure 2.**
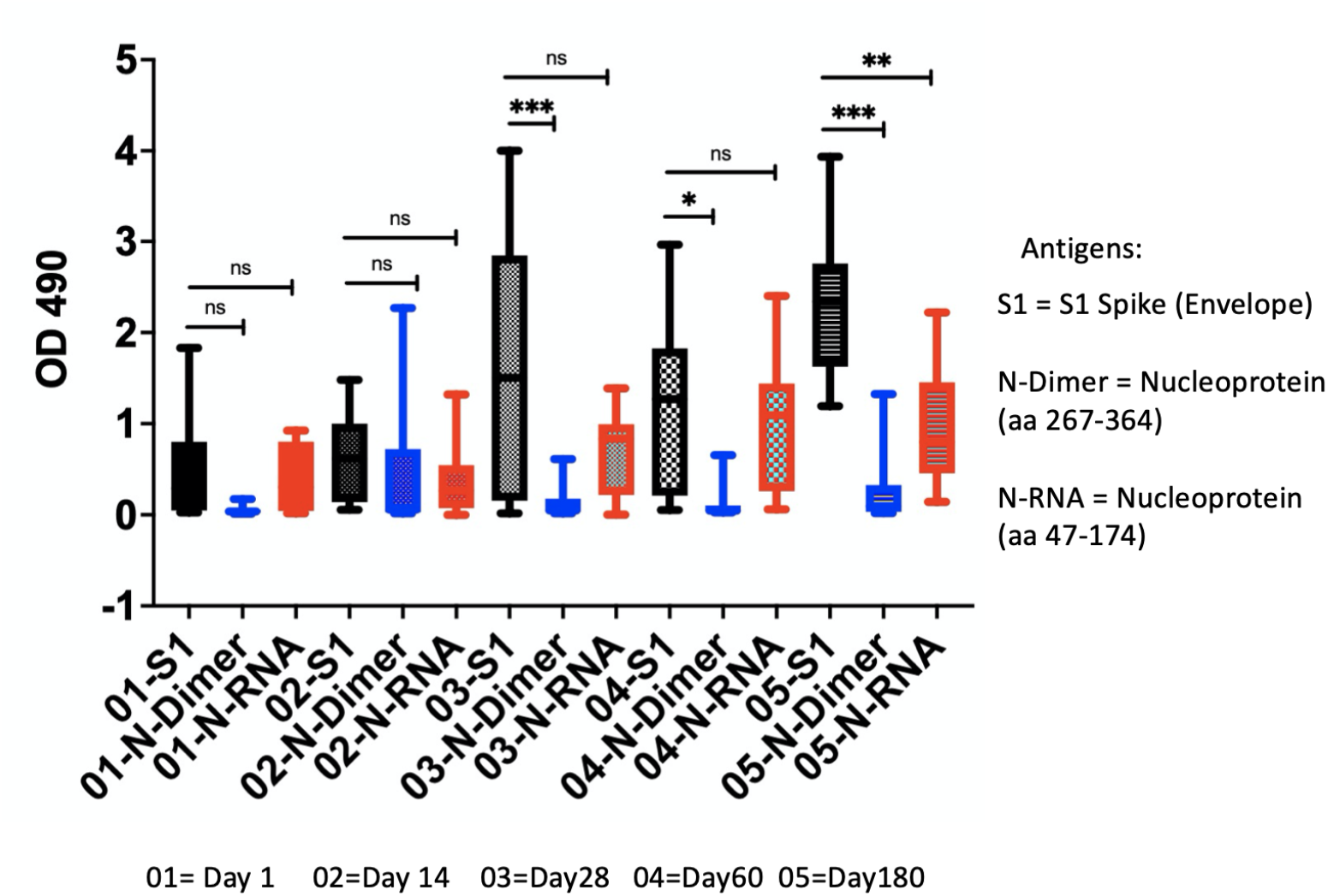
Comparison of serum IgG reactivity in thirty-two Covid-19 patients with recombinant S1 Spike protein, the two recombinant nucleoproteins (aa47-174/NC-RNA and 247-364/NC-Dimer) during a 6-month follow-up period. Significantly higher serum IgG reactivity was seen against the S1-spike and the NC-RNA aa47-174-nucleoproteins. Boxplots showing mean reactivity and range per timepoint is shown.

A subgroup of twenty-two individuals was tested for nasopharyngeal swab collected IgA reactivity and positive reactivity was seen in upto thirty-six percent of tested individuals at least once between days 14 to 60 against the S1 Spike protein (Figure 3).

**Figure 3.**
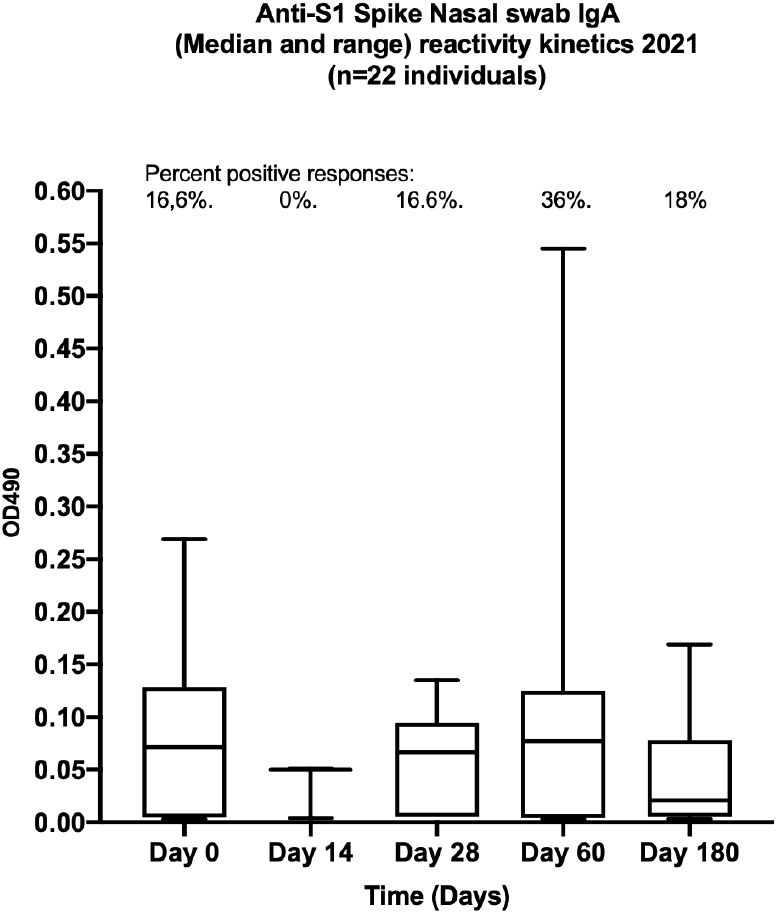
Nasal swab IgA ELISA reactivity seen in twenty-two Covid-19 patients against the SARS-CoV-2 surface recombinant S1 Spike-protein. Boxplots showing median reactivity and range per timepoint is shown.

## Discussion

Serological analysis directed against the principal SARS-CoV-2 neutralizing protein the viral S1 Spike protein has been presented to be one of the most long-lasting immune responses in previous studies (11, 13). In this study all involved patients developed detectable anti-S1-binding IgG and IgA responses at least once during the study period. Ninety-six percent of the study participants were seropositive at 6 months. All showed presence of both serum IgG (IgG1) and IgA, and in a subgroup also local mucosal nasopharyngeal sIgA against the outer surface proteins of the SARS-CoV-2 for upto 2 months follow-up.

It is of importance that the serum IgG against neutralizing viral proteins remain elevated, and hopefully at a protective level, even though no clear antibody-titer or antibody quantity limit on correlation from infection or reduced transmissibility have been determined yet. Similarly, it was valuable to see that also serum IgG against the nucleoprotein was frequent among the infected patients. However, serum IgG/IgA against the NC-proteins have been described as both negative and positive for human health as described in different patient categories. In one study higher amounts of these immunoglobulins correlate with antibody-dependent enhancement (ADE) of infection, even though no clear mechanistic process was presented (14), and in other studies these antibodies have been part of the cure/health promoting immune response. The possible positive interpretations or explanations given would be the clearance of circulating viral proteins that otherwise may cause formation of immune complexes, or result in inflammation at local sites where the NC-proteins aggregate (15).

It is estimated that around half of hospital admissions for young children and around twenty percent of hospitalizations of pneumonias among elderly is due to respiratory viral infections, but before 2020 mainly due to respiratory syncytial virus or influenza viruses (16). From 2020, these infectious agents have suddenly become considerably less frequent, and exchanged to the completely dominating betacoronavirus SARS-CoV-2 (in Scandinavia). Respiratory infectious viruses (historically dominated by around eighty to ten different virus families will thus remain serious health challenges even though decades of intensive research for developing antiviral drugs and vaccines against them have been performed.

Still, it is important to identify the correlates of protective immunity, and sofar the best and most often defined correlate in almost any respiratory infectious illness have been antibodies (17). Antibodies have been shown to neutralize infectivity by attaching to viral surface proteins, and in addition to neutralization to trigger complement-activation and provide antibody-dependent cell-mediated cytotoxicity against infected viruses and cells. There are few reasons yet to believe that the SARS-CoV-2 will not be sensitive to the same correlates of protection (18, 19, 20). The challenge will be to obtain antibodies capable of recognizing the new circulating S1-envelope protein mutant SARS-CoV-2 strains, especially close to the nasal mucosal surfaces as seen in a subpopulation (Fig.3) here. The possibly positive result with obtaining the SARS-CoV-2 specific immunity through natural infection may be the broader immune responses towards several viral proteins, several that are not included in the vaccines. Immune memory B- and T-lymphocytes will thus be obtained both against the viral surface proteins and several hopefully more conserved intraviral proteins such as the nucleo-proteins (N) and the non-structural proteins (nsp:s).

The question is just to understand the longevity and the magnitude of these immune responses in Covid-19 patients with mild and/or severe disease as well as in the SARS-CoV-2 vaccinated individuals of different age groups and genders.

The conclusions from this study were that the majority of Covid-19 patients develop a 6-months lasting humoral serum and mucosal immunity towards the outer S1 spike protein of SARS-CoV-2.

## ACKNOWLEDGEMENTS

We are most grateful for the invaluable support with SARS-CoV-2 reagents to Dr. Ackloo S. SGU Toronto, ON, Canada. All involved researchers confirm no competing interests.

## References

1. Böttiger M. Polio immunity to killed: an 18-year follow-up. Vaccine 1990, 8(5):443–5. Doi: 10.1016/0264-410x(90)90244-g.

2. Böttiger M, Larsson B. Swedish inactivated polio vaccine: laboratory standardization and clinical experience over a 30-year period. Biologicals 1992, 20(4):267–75. Doi: 10.1016/s1045-1056(05)80046-6.

3. Haralambieva IH, Kennedy RB, Ovzyannikova IG, Schaid DJ, Poland GA. Current perspectives in assessing humoral immunity after measles vaccination. Expert Rev.Vaccines. 2019, 18(1):75–87. Doi: 10.1080/14760584.2019.1559063.

4. Mörner A, Bråve A, Kling AM, Kuhlmann-Berenzon S, Krook K, Hedenskog M, Silhammar I, Ljungman M, Örtqvist A, Andresson S, Brytting M, Thorstensson R, Linde A. Pandemic influenza A(H1N1)pdm09 seroprevalence in Sweden before and after the pandemic and the vaccination campaign in 2009. PLoS One 2012, 7(12):e53511. Doi: 10.1371/journal.pone.0053511.

5. Truelove S, Zhu H, Lessler J, Riley S, Read JM, Wang S, Kwok KO, Guan Y, Jiang CQ, Cummings DA. A comparison of hemagglutination inhibition and neutralization assays for characterizing immunity to seasonal influenza A. Influenza Other Respir Viruses, 2016, 10(6):518–524. Doi: 10.111/urv.12408.

6. Buchwald AG, Graham BS, Traore A, Haidara FC, Chen M, Morabito K, Lin BC, Sow SO, Levine MM, Pasetti MF, Tapia MD. RSV neutralizing antibodies at birth predicts protection from RSV illness in infants in the first three months of life. Clin Infect Dis. 2020, ciaa648. Doi: 10.1093/cid/ciaa648.

7. Zost SJ, Gilchuk P, Case JB, Binshtein E, Chen RE, Nkolola JP, Schäfer A, Reidy JX, Trivette A et al. Potently neutralizing and protective human antibodies against SARS-CoV-2. Nature 2020, 584(7821):443–449. Doi: 10.1038/s41586-020-2548-6.

8. Zhou G, Zhao Q. Perspectives on therapeutic neutralizing antibodies against the Novel Coronavirus SARS-CoV-2. J Biol Sci. 2020, 15;16(10):1718–1723. Doi: 10.7150/ijbs.45123.

9. Rudberg AS, Havervall S, Månberg A, Jernbom Falk A, Aguilera K, Ng H, Gabrielsson L, Salomonsson AC, Hanke L, Murrell B, McInnerney G, Olofsson J, Andersson E, Hellström C, Bayati S, Bergström S, Pin E, Sjöberg R, Tegel H, Hedhammar M, Philipsson M, Nilsson P, Haber S, Thålin C. SARS-CoV-2 exposure, symptoms and seroprevalence in healthcare workers in Sweden. Nat. Commun. 2020, 11(1):5064. Doi: 10.1038/s41467-020-18848-0.

10. Varnaite R, Garcia M, Glans H, Maleki KT, Sandberg JT, Tynell J, Christ W, Lagerqvist N, Asgeirsson H, Ljunggren HG, Ahlen G, Frelin L, Sällberg M, Blom K, Klingström J, Gredmark-Russ S. Expension of SARS-CoV-2-specific antibody-secreting cells and generation of neutralizing antibodies in hospitalized COVID-19 patients. J.Immunol. 2020, 205(9):2437–2446. Doi: 10.4049/jimmunol.2000717.

11. Marklund E, Leach S, Axelsson H, Nyström K, Norder H, Bemark M, Angeletti D, Lundgren A, Nilsson S, Andersson LM, Ylimaz A, Lindh M, Liljeqvist JÅ, Gisslen M. Serum-IgG responses to SARS-CoV-2 after mild and severe COVID-19 infection and analysis of IgG non-responders. PLoS One 2020, 15(10):e241104. Doi: 10.1371/journal.pone.0241104.

12. Strålin K, Wahlström E, Walther S, Bennet-Bark AM, Heurgren M, Linden T, Holm J, Hanberger H. Second wave mortality, among patients hospitalized for COVID-19 in Sweden: a nationwide observational cohort study. medRxiv https://doi.org/10.1101/2021.03.29.21254557

13. Chiereghin A, Zagari EM, Galli S, Moroni A, Gabrielli L, Venturoli S, Bon I et al. Recent advances in the evaluation of serological assays for the diagnosis of SARS-CoV-2 infection and COVID-19. Front Public Health. 2021, 8:620222. Doi:10.3389/fpubh.2020.620222.

14. Batra M, Tian R, Zhang C, Clarence E, Sacher CS, Miranda JN et al. Role of IgG against N-protein of SARS-CoV2 in CIVID19 clinical outcomes. Sci Rep. 2021, 11:3455, doi.org/10.1038/s41958-021-83108-0.

15. Roncati L. Type 3 hypersensitivity in Covid-19 vasculitis. Clin.Immunol. May 26, 2020, doi.org/10.1016/j.clim.2020.108487.

16. Ruuskanen O, Lahti E, Jennings LC, Murdoch DR. Viral pneumonia. Lancet 2011, 377, 1264–1275.

17. Plotkin SA, Plotkin SA. Correlates of vaccine-induced immunity. Clin. Inferct. Dis. 2008, 47:401–409.

18. O’Nions J, Muir L, Zheng J, Rees-Spear C, Rosa A, Roustan C, Earl C, Cherepanov P, Gupta R, Khwaja A, Jolly C, McCoy LE. SARS-CoV-2 antibody responses in patients with acute leukemia. Leukemia 2021, 35:289–292.

19. Ng KW, Faulkner N, Cornish G, Rosa A, Hussasin S, Harvey R et al. Pre-existing and de novo humoral immunity to SARS-CoV-2 in humans. Science 2020. http://doi.org/10.1126/scienceabe1107.

20. Seow J, Graham C, Merrick B, Acors S, Pickering S, Steel KJA et al. Longitudinal observation and decline of neutralizing antibody responses in the three months following SARS-CoV-2 human infection. Nature Microb. 2020. http://doi.org/10.1038/s41564-020-00813-8.

